# A pangenome analysis pipeline (PSVCP) provides insights into rice functional gene identification

**DOI:** 10.1101/2022.06.15.496234

**Authors:** Jian Wang, Wu Yang, Shaohong Zhang, Haifei Hu, Yuxuan Yuan, Jingfang Dong, Luo Chen, Yamei Ma, Tifeng Yang, Lian Zhou, Jiansong Chen, Bin Liu, Chengdao Li, David Edwards, Junliang Zhao

## Abstract

**Background:** A pangenome aims to capture the complete genetic diversity within a species and reduce bias in genetic analysis inherent in using a single reference genome. However, the current linear format of most plant pangenomes limits the presentation of position information for novel sequences. Graph pangenomes have been developed to overcome this limitation. However, there is a lack of bioinformatics analysis tools for graph format genomes.

**Results:** To overcome this problem, we have developed a novel pangenome construction strategy and a downstream pangenome analysis pipeline that captures position information while maintaining a linearized layout. We applied this strategy to construct a high-quality rice pangenome using 12 representative rice genomes and analyze an international rice panel with 413 diverse accessions using the pangenome reference. Our results provide insights into rice population structure and genomic diversity. Applying the pangenome for PAV-based GWAS analysis can identify causal structural variations for rice grain weight and plant height, while SNP-based GWAS can only identify approximate genomic locations. Additionally, a new locus (qPH8-1) was found to be associated with plant height on chromosome 8 that could not be detected using the SNP-based GWAS.

**Conclusions:** Our results demonstrate that the pangenome constructed by our pipeline combined with PAV-based GWAS can provide additional power for genomic and genetic analysis. The pangenome constructed in this study and associated genome sequence data provide valuable genomic resources for future rice crop improvement.

## Background

Rice (*Oryza sativa L*) is one of the most important staple crops, feeding nearly half of the world’s population. As this population expands to 10 billion people, there is an urgent need to increase the productivity of crops, while facing the impact of climate change on agricultural productivity. The application of genomics assisted breeding is seen as one of the best opportunities to increase crop productivity, with the exploitation of diversity stored in germplasm collections as a major resource for crop improvement [1]. With rapid advances in DNA sequencing technologies, genomic diversity within rice germplasm has been characterized by resequencing thousands of individuals and comparing the resulting data with reference genome assemblies. However, it is now understood that a single reference genome does not represent the genomic diversity of a species due to significant sequence presence/absence variation (PAV) between individuals [2]. To capture the genomic variations in a population, pangenome assemblies have been constructed. Pangenomes represent the gene content of a species rather than a single individual [3], and using a pangenome as a reference, structure variations (SVs) can be more easily and accurately genotyped by low cost short-read sequencing technologies, facilitating efficiently characterisation of genomic diversity within a species.

Pangenomes have now been constructed and analyzed for several crop species, including wheat, Brassicas, barley, banana and pigeonpea [4-8]. Several pangenomes have been constructed in rice, and pangenomic analyses have identified genome sequences that are absent in the Nipponbare reference, the most commonly used reference in rice genomic studies [9-11]. For example, a study using 3,010 rice accessions identified 268 Mb of new sequences, with 12,465 new genes, and 19,721 dispensable genes compared to the Nipponbare reference genome [12].

Recent advances in pangenomics have led to the construction of graph-based pangenomes [13, 14] that code genetic variants as nodes and edges, and preserve the contiguity of the sequence and structural variation between individuals [15]. Graph-based pangenome approaches are relatively new, but have been applied to important crops, including soybean, bread wheat, and rice [10, 16-18]. Though graph based pangenomes have advantages, they also suffer limitations; for example, as most genome analysis tools were developed for linear sequences, scalable software and mature data structures suitable for graph-based pangenome analysis are still limited. A linear format pangenome with a fixed order coordinate system is still valuable for genomic studies, however, they struggle to represent the position of SVs and so potentially lose valuable information.

In this study, we developed a pangenome construction strategy that can preserve SV position, embedding them into a linear pangenome. We also developed a suite of tools for mapping short sequencing reads to this pangenome for PAV genotyping that can recover the genomic position of sequence variations. We applied this pipeline to construct a rice pangenome using 12 diverse accessions representing major subpopulations of Asian rice and identify PAVs from an international rice mini core panel of 413 accessions [19]. This revealed extensive genomic diversity among rice germplasm, and PAV-based population analysis provided insights into population structure and successfully identified causal PAVs that impact grain weight and plant height. This study presents a new tool for pangenome analysis and provides valuable genomic resources for rice functional genomics, demonstrating the advantages of using a coordinate linked linear pangenome to identify PAVs for functional analysis.

## Results

### A novel pangenome construction and PAV analysis pipeline

In this study, we developed a pangenome construction and PAV genotype calling pipeline (PSVCP) (Additional file 1: Fig. S1). The pipeline includes three main steps, 1) Iterative alignment between genomes to identify novel segments, then the integration of these sequences into the reference genome to construct a pangenome (Fig. 1A). 2) Mapping of short-read resequencing data to the pangenome to detect PAVs based on read coverage (Fig. 1B). 3) Calling PAV genotypes at the population level based PAVs from all accessions’ (Fig. 1C).

**Fig 1.**
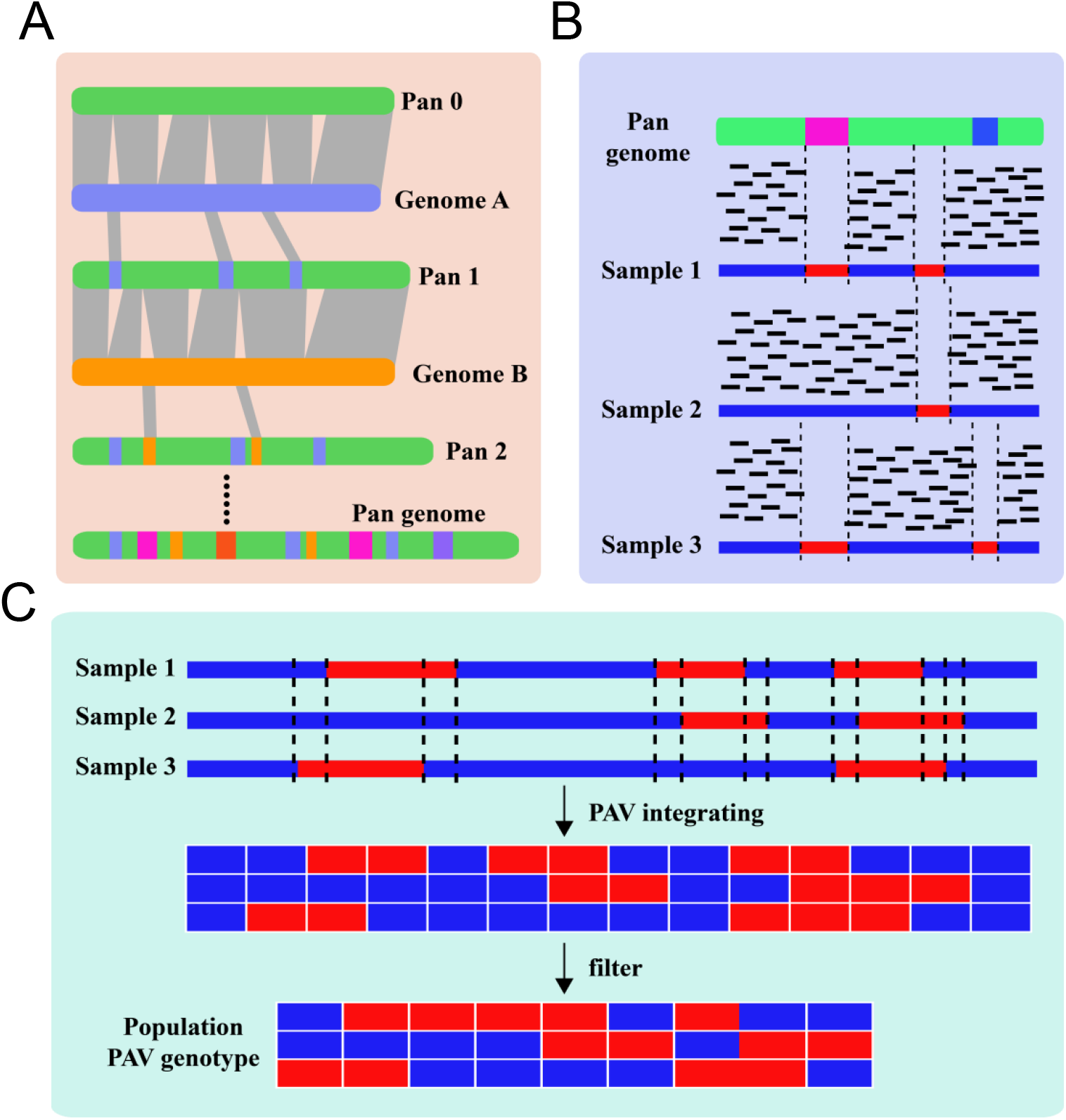
Scheme diagram of PSVCP pipeline. **A** construction of linearized pan-genome. **B** PAV was re-calling by sequencing coverage calculation. **C**. population PAV genotype calling

We initially selected 12 assembled genome sequences of cultivated rice, including 11 Asian cultivated rice (*O. sativa*) accessions selected from 33 representative accessions based on their subpopulation [10] and one African cultivated rice (*Oryza glaberrima*) (Additional file 2: Table S1) for pangenome construction using Nipponbare as the primary reference [20]. A total of 24,585 novel sequences were identified and inserted into the Nipponbare reference. The mean, median, maximum and the sum of insertion lengths are 2,607 bp, 338bp, 96,797 bp, and 64.10 Mbp respectively (Additional file 1: Fig. S2A, B). A subset of these sequences was validated by amplification and sequencing (Additional file 1: Fig. S3).

We analyzed the distribution of these additional sequences and found that 43.1% overlapped ±2 kb upstream/downstream of genes, while 35.7% overlapped with genic regions (Additional file 1: Fig. S2C). Altogether, 6,797 sequences were inserted into 5,925 Nipponbare genes (Fig. 2). A total of 1,939 new genes were de novo annotated, and functional analysis suggests that they are enriched with terms associated with photosynthesis, the generation of precursor metabolites and energy (Additional file 2: Table S2). Modelling suggests that the initial 12 rice accessions were sufficient to capture the majority of sequence diversity within rice (Additional file 1: Fig. S4).

**Fig. 2.**
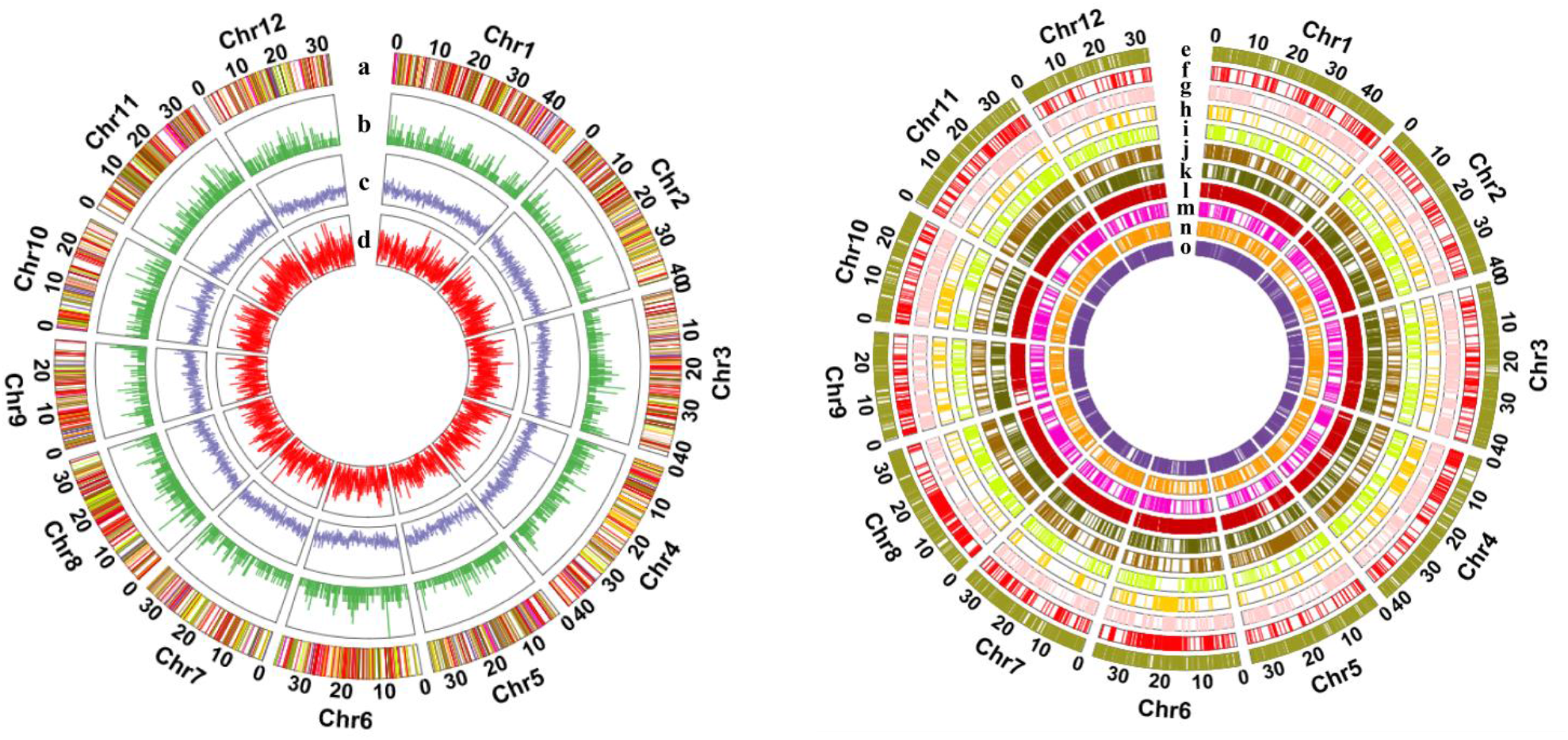
Linear pan-genome constructed by 12 rice accession a. New genes transported with PAV b. Genes in MSU interrupted by PAV c. Pan genes density d. PAV density e. PAV from CG14.fa; f. PAV from Basmati1.fa; g. PAV from N22.fa; h. PAV from FH838.fa; i. PAV from Tumba.fa; j. PAV from TM.fa; k. PAV from R498.fa; l.PAV from CN1.fa; m. PAV from LJ.fa; n. PAV from NamRoo.fa; o. PAV from Lemont.fa

The completeness of the pangenome was evaluated using Benchmarking Universal Single-Copy Orthologs (BUSCO) [21] (Additional file 2: Table S3). Of the 1614 single-copy orthologs identified in embryophytes, 98.8% were complete in our assembly, which is similar to or a little higher than the 3K rice pangenome (98.5%) [12] (Additional file 2: Table S3). We mapped resequencing data for 413 rice accessions collected from a diverse international panel (RPD2) [19] to the pangenome and the Nipponbare genome respectively. The results showed the average mapping rate to the pangenome was 97.84%, which is higher than the mapping rate to the Nipponbare reference (93.05%) (Additional file 1: Fig. S5A). These results demonstrate that our pangenome captured more diversity than the single Nipponbare reference.

### Population-wide TE and PAV analysis in an international diverse rice panel

Illumina whole-genome sequencing data was generated for 413 accessions representing an international rice collection from 96 countries [19]. The reads were mapped to the pangenome and PAVs were genotyped using the PSVCP pipeline. This identified an average of 99,239 PAVs (>50 bp) per accession, ranging from 38,052 to 213,931. Around 85% of the inserted sequences were transposable elements, with 40% annotated as Gypsy LTR-retrotransposons and 28.6% as Helitron DNA transposons (Additional file 2: Table S4). We examined the diversity of representative retrotransposon families across all 413 accessions [22]. In total, 66,441 variable retrotransposon sequences were identified, with 29,281 (44%) absent from the Nipponbare reference assembly.

Retrotransposon abundance ranged from 12 (Rn60/Gypsy) to 15,599 copies (Rire3 /Gypsy). Notably, half of the copies in the retrotransposon TE families Rn60, Rire3, Fam81-fam82, Rire2, Hopi, Fam93_ors14, Fam51_osr4 and Tos17 were not identified in the Nipponbare reference. The majority of retrotransposons were from Hopi, Fam81-fam82 and Rire3 TE families, which belong to the Gypsy family, and most of these originate from *Indica* accessions, suggesting an expansion of Gypsy elements in *Indica* compared to *Japonica* [23, 24]. TE families Fam93_ors14, Hopi and Fam81-fam82 show significantly higher frequency in *Indica* than *Japonica* and *Aus* accessions, while the Rire3 family is less abundant in *Aus* varieties compared to the other populations (Additional file 2: Table S5). This suggests ongoing transposition during domestication and subsequent breeding.

We identified 11,617 (28.9%) dispensable genes across the 413 rice accessions (Additional file 2: Table S6). Annotation suggests that these are enriched for functions associated with protein phosphorylation, telomere maintenance, DNA duplex unwinding, photosynthesis, defence response and pathogenesis (Additional file 2: Table S7), which is similar to the findings in other crop pangenome studies [25, 26]. We observed a significant difference in average gene numbers between *Japonica, Indica and Aus* (Fig. 3A). *Japonica* contains the most genes (48,884 ± 472), with fewer genes in *Indica* (47,455 ± 537) and *Aus* (47,441 ± 405). The difference in average gene number hides a complex pattern of increases and decreases in the frequency of specific genes (Fig. 3B). A total of 978 genes show increased frequency in *Japonica*, while 2,986 genes show decreased frequency. Genes showing increased frequency are enriched in functions associated with DNA integration (Additional file 2: Table S8), while genes showing decreased frequency are annotated with disease resistance terms, including pathogenesis and defence response (Additional file 2: Table S9). Among the 2,986 genes with lower frequency in *Indica*, 116 (3.8%) genes are absent from the Nipponbare reference. In contrast, of the 978 genes exhibiting higher frequency in *Indica*, 513 (52.5%) genes are absent from the Nipponbare reference, with 482 derived from the *Indica* rice genomes. This reflects differences in gene content between sub-species at the population level.

**Fig. 3.**
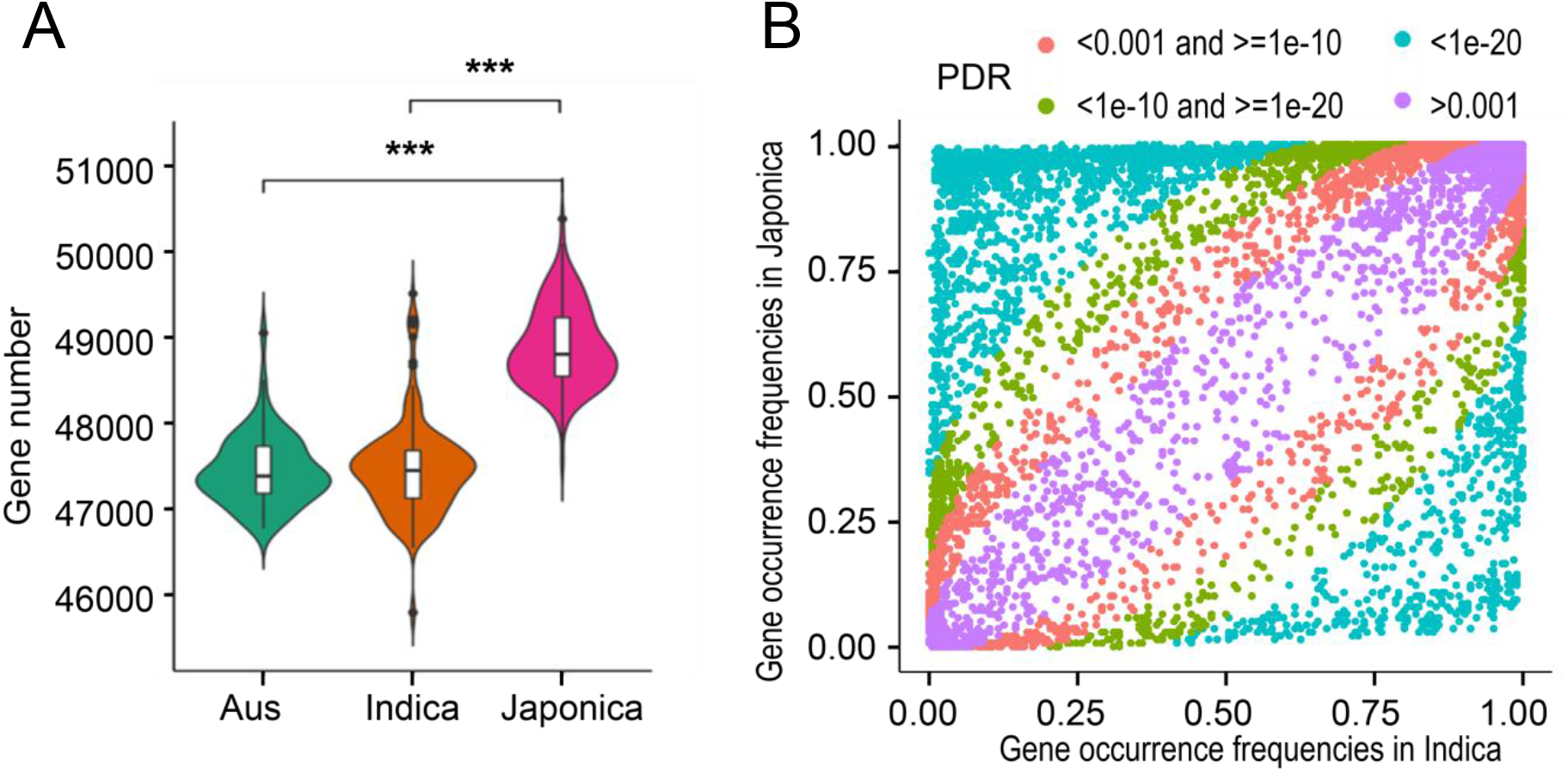
Gene number and frequency analysis among Subpopulation. A Violin plots showing gene abundance for the *Aus*, In*dica* and *Japonica* significance differences between groups are indicated (***p < .005). **B** Comparison of gene frequency between *Indica* and *Japonica*. Colors indicate p-value

### Population structure analysis based on pangenome PAVs

We performed population genetic analysis in the international panel using PAVs and compared the results with SNP-based analysis. The mean fixation index (Fst) between populations estimated using the SNP data (*Japonica*-*Indica*: 0.476 ±0.207, *Japonica-Aus*: 0.525 ±0.205 and *Indica-Aus*: 0.304 ±0.158) is higher than calculated using PAV data (*Japonica-Indica*: 0.416 ±0.183, *Japonica-Aus*: 0.430 ±0.184 and *India-Aus*: 0.204 ±0.128) (Additional file 2: Table S10). Fst analysis results show similar distribution trends between PAVs and SNPs on the whole genome scale (Additional file 1: Fig. S6). SNP-based analysis shared Fst differentiation regions with PAV-based analysis (within the top 1% Fst windows) between populations. For example, both SNP and PAV results share 33 out of 54 of the *Japonica*-*Indica* Fst differentiation regions, which contained 376 genes. We analysed 15 well-studied domestication and improvement associated genes to compare the Fst detection between SNP and PAVs. Among the 15 genes, three were within the top 10% of FST differentiation regions among *Indica, Japonica* and *Aus* subpopulations using SNP and PAV data (Additional file 2: Table S11). We also detected regions displaying significant differences between Fst values based on PAVs and SNPs. To investigate this further, we selected a prominent region at 7.2-9.2 Mbp of chromosome 8 where we observed a much higher Fst value between *Indica* and *Japonica* calculated by PAVs than SNPs (Fig. 4A). Further analysis revealed that PAVs could detect more genetic diversity than SNPs in this region (Fig. 4A). The region showed a higher ratio of novel sequences than the Nipponbare reference. The length of this region is about 1,600 kb in Nipponbare, while in the pangenome, the interval is 2 Mb, with 271 annotated genes, of which 162 are transposons.

**Fig 4.**
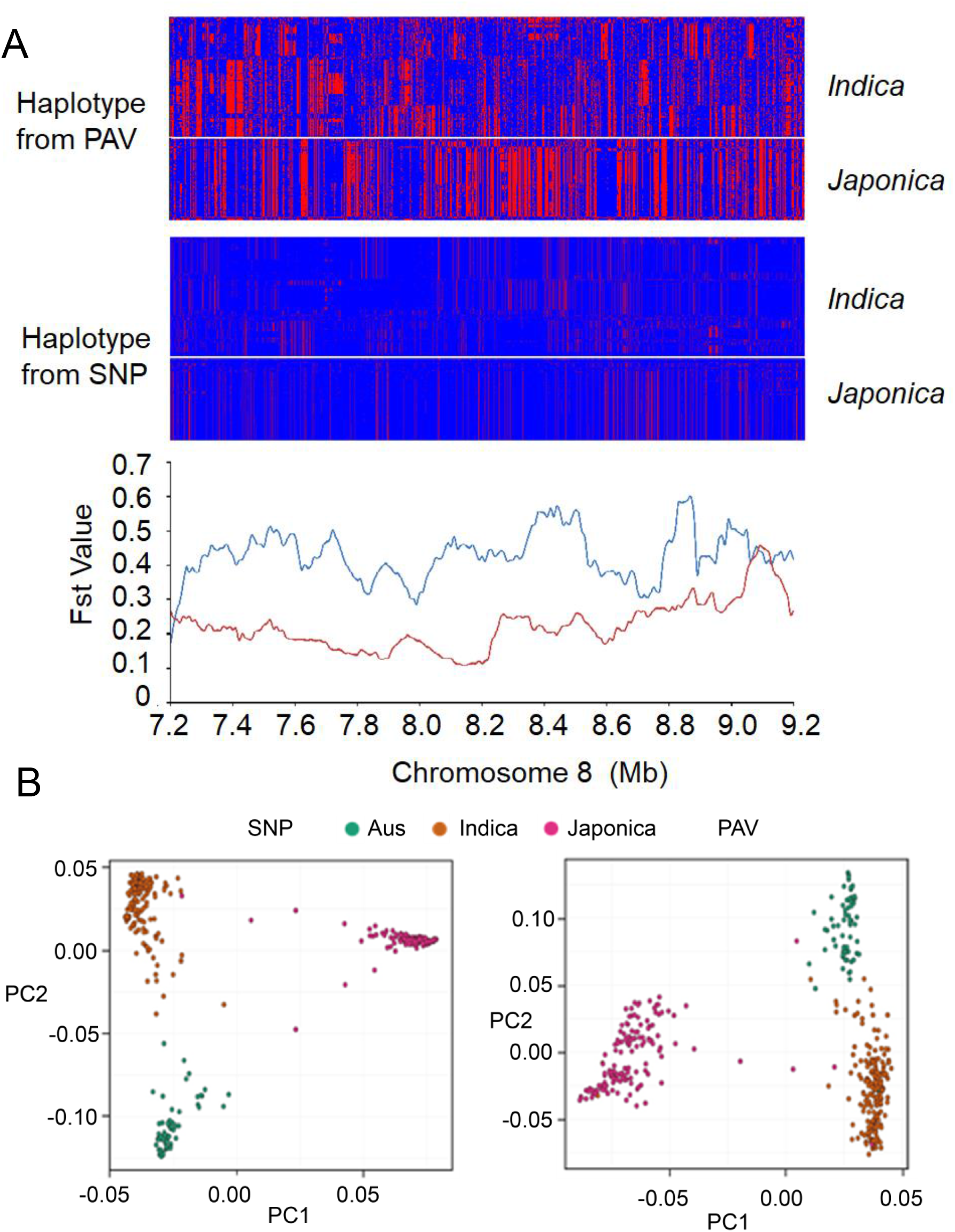
Population structure analysis based on PAV and SNP A haplotype pattern and Fst analysis by PAV and SNP datga in 7.2-9.2 Mb of chromosome 8 in the rice pangenome. **B** PCA plot generated by PAV and SNP data

PAV-based population structure shows similar clustering to SNP-based phylogeny, with 413 accessions clustered into three main subpopulations. However, the PAV-based phylogeny does not cluster individuals completely according to subpopulations, and the PAV-based PCA suggests a greater variation between rice accessions than the SNP-based analyses (Fig. 4B). For example, accessions in *Indica* and *Aus* subpopulations were grouped into two clusters compared with the SNP-based PCA result, and some accessions in the *Indica* subpopulation clustered with the *Aus* subpopulation. A similar pattern was observed in the PAV-based phylogeny with 73 *Indica* accessions clustering with the *Aus* subpopulation (Additional file 1: Fig. S7).

### Using pangenome to perform PAV-based GWAS

As a pangenome permits the genotyping of a greater amount of genetic diversity than a single reference, it supports more powerful genetic analysis, capturing missing heritability. To explore this additional potential, particularly for identifying functional PAVs underlying QTLs, we conducted GWAS for two important agronomic traits of rice, thousand grain weight (TGW) and plant height (PH), using SNPs genotyped from Nipponbare and PAVs genotyped across the pangenome.

For TGW, the SNP-GWAS identified 354 significant associations (Additional file 1: Fig. S8A), with the most significant located in Nip Chr5: 5,375,764 bp (pangenome Chr5: 6,017,339 bp), 9,063 bp away from *GW5*, a known functional gene controlling rice grain weight [26]. However, none of the associated SNPs were the causal variations of *GW5*, which are two PAVs (950-bp and 1,212-bp) in the promoter region, controlling the grain weight phenotype [27]. Our pangenome can capture these PAVs, which are absent in the Nipponbare reference genome. Using the pangenome, PAV-GWAS narrowed down the association signal in the same interval as SNP-GWAS (Fig. 5A; Additional file 1: S8A) and also detected the most significant associated signal as the causal variations of GW5 (Fig. 5B, C). We further analyzed the PAV genotypes and identified three haplotypes. The accessions with Hap1 (with both 1,212 bp and 950 bp PAVs) showed significantly lower grain weight than accessions with the other two haplotypes (Hap2, Hap3) with p-values (two-tailed student’s t-test) of 3×10^−5^ and 3×10^−9^ respectively (Fig. 5C). This result is in accord with a previous study that demonstrated that the 950 bp deletion decreased the expression of the functional gene (*qSW5*), while the 1,212 bp deletion disrupts the coding region. Both deletions will lead to grain width and weight phenotype variations [26].

**Fig. 5.**
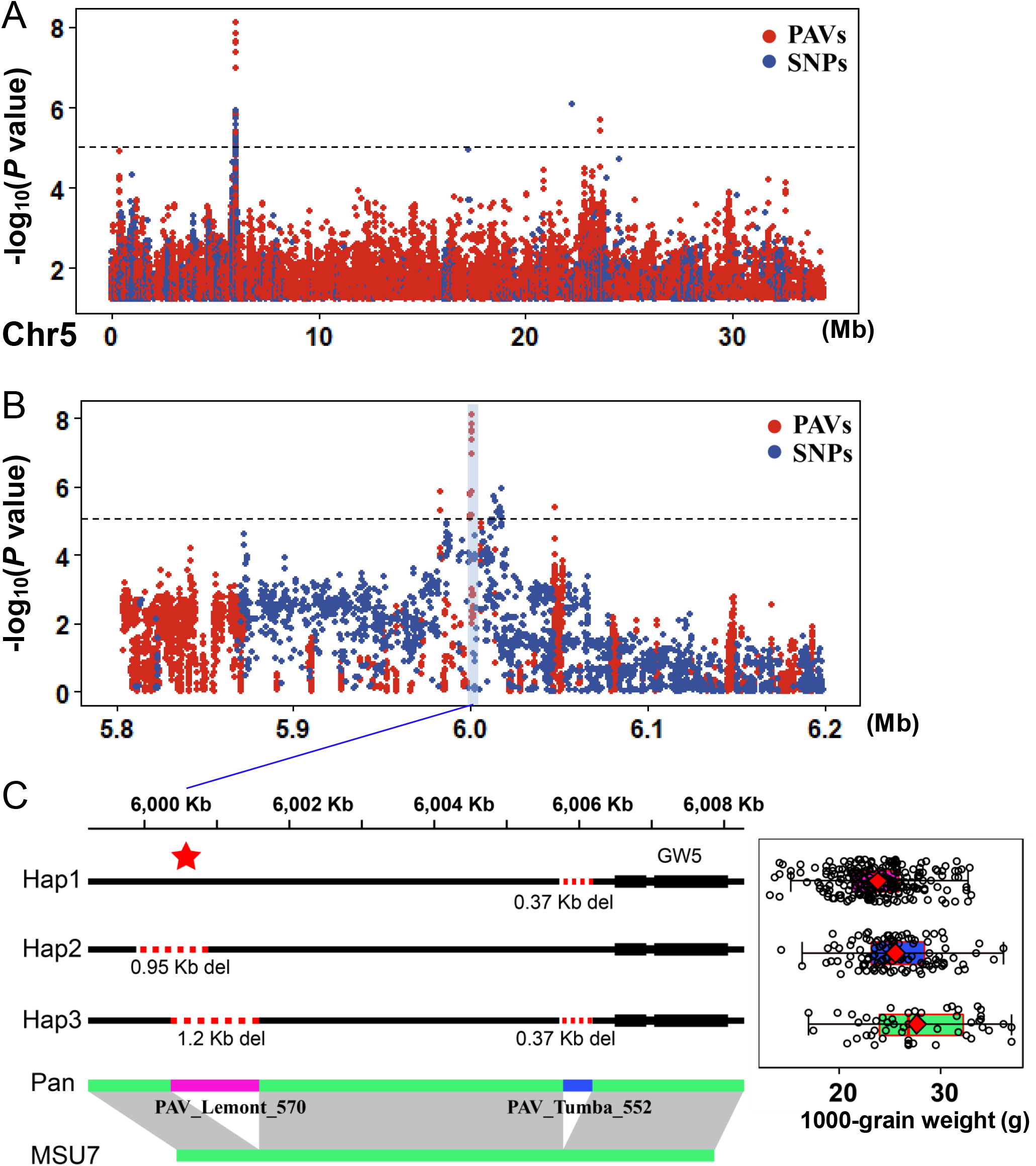
GWAS of thousand grain weight in 413 accessions population. **A**,**B** Manhattan plots for thousand grain weight analysed by SNP-GWAS and PAV-GWAS. **C** Haplotype analysis in *GW5* promoter region

SNP-GWAS identified 37 SNPs associated with plant height (Additional file 1: Fig. S8B). Similar to the TGW GWAS result, both SNPs and PAV-GWAS were able to locate previously characterized locus harboring the “Green Revolution Gene” (*sd1*) [28]. The most significant PAV is located inside the *sd1* gene, a previously reported causal variation determining plant height in rice (Additional file 1: Fig. S9) [28]. Statistical analysis shows that this PAV is significantly correlated with the PH phenotype (two-tailed student’s t-test, p-value: 3.3 × 10^−29^), further validating the accuracy of PAV-GWAS. Furthermore, we also identified a novel locus (*qPH8-1*) controlling PH in rice on chromosome 8 by PAV-GWAS (interval: 4,660,000-4,860,000 bp in the pangenome), that was not identified by SNP-GWAS (Fig. 6). The most significant PAV was a 13 kb sequence containing two retrotransposon genes (*LOC_Os08g07410, LOC_Os08g07420*) located 1 kb upstream of *LOC_Os08g07400*. This sequence was present in 288 out of the 413 accessions, and the accessions without the 13 kb sequence had significantly greater plant height (two-tailed student’s t-test, p-value: 5.7 × 10^−20^) than those had the 13 kb sequence. Expression analysis shows that the presence or absence of this 13 kb sequence is significantly correlated with the expression level of *LOC_Os08g07400*, which is located 2 kb downstream from the PAV (Fig. 6C). These results suggest that this PAV, caused by retrotransposon movement, may impact downstream gene expression and plant height phenotype. The mechanisms underlying the discordance of results between SNP-GWAS and PAV-GWAS in this PH QTL were further investigated. We examined the genome structure landscape at the population level and examined the relationship between the 13 kb PAV and the nearby SNPs. The presence or absence of the 13 kb sequence strongly correlates with the plant height phenotype (Fig. 7A). However, the SNPs on both sides of the PAV did not associate with the plant height. Linkage disequilibrium (LD) analysis further demonstrated the PAV interval formed an LD block, while the PAV genotype did not correlate with the SNP phenotype (Fig. 7B).

**Fig. 6.**
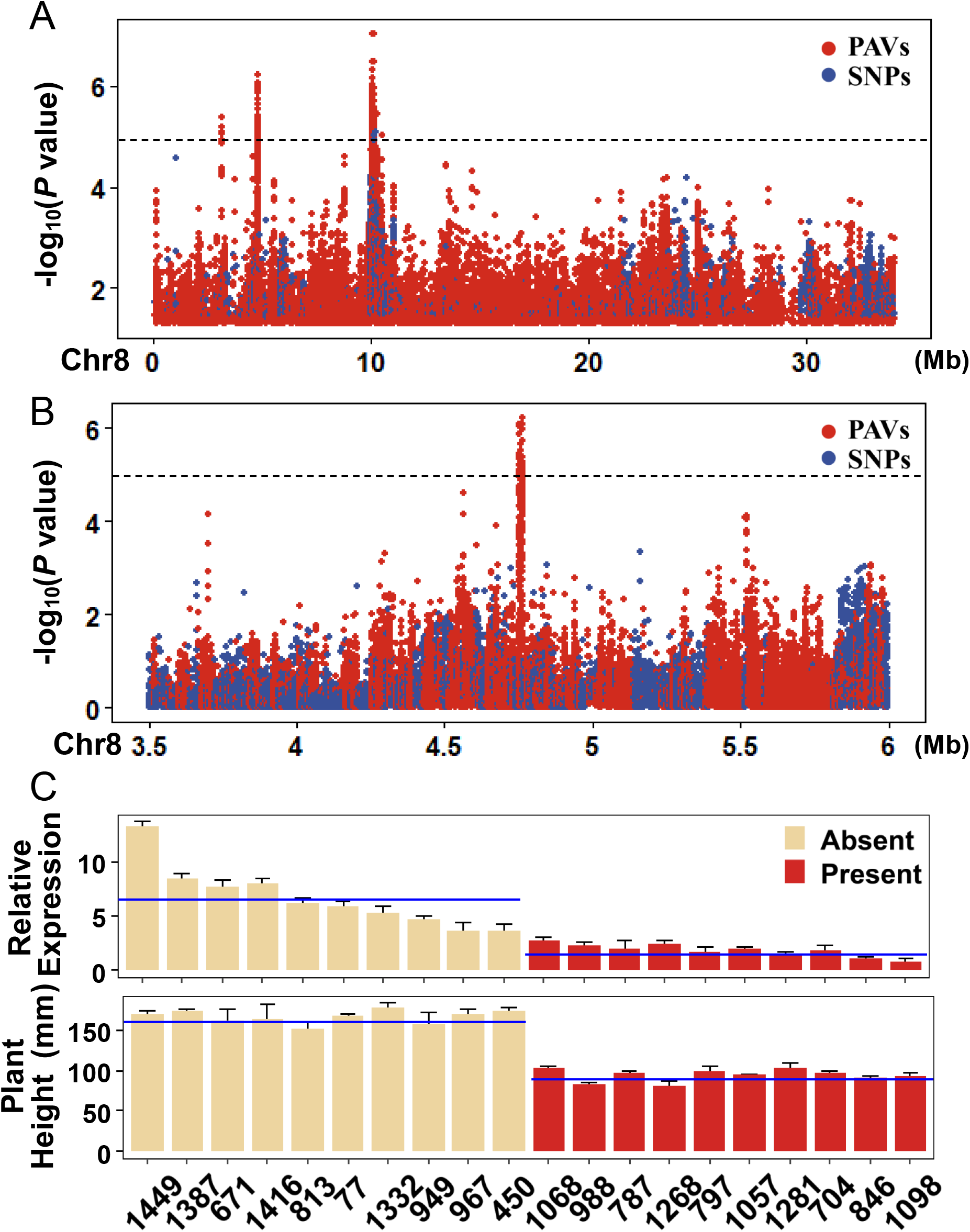
GWAS of plant height in 413 accessions population. **A**,**B** Manhattan plots for plant height analysed by SNP-GWAS and PAV-GWAS. **C** Expression analysis of *LOC_Os08g07400* in the accessions with absence and presence of 13-kb. The blue line is the mean value.

**Fig. 7.**
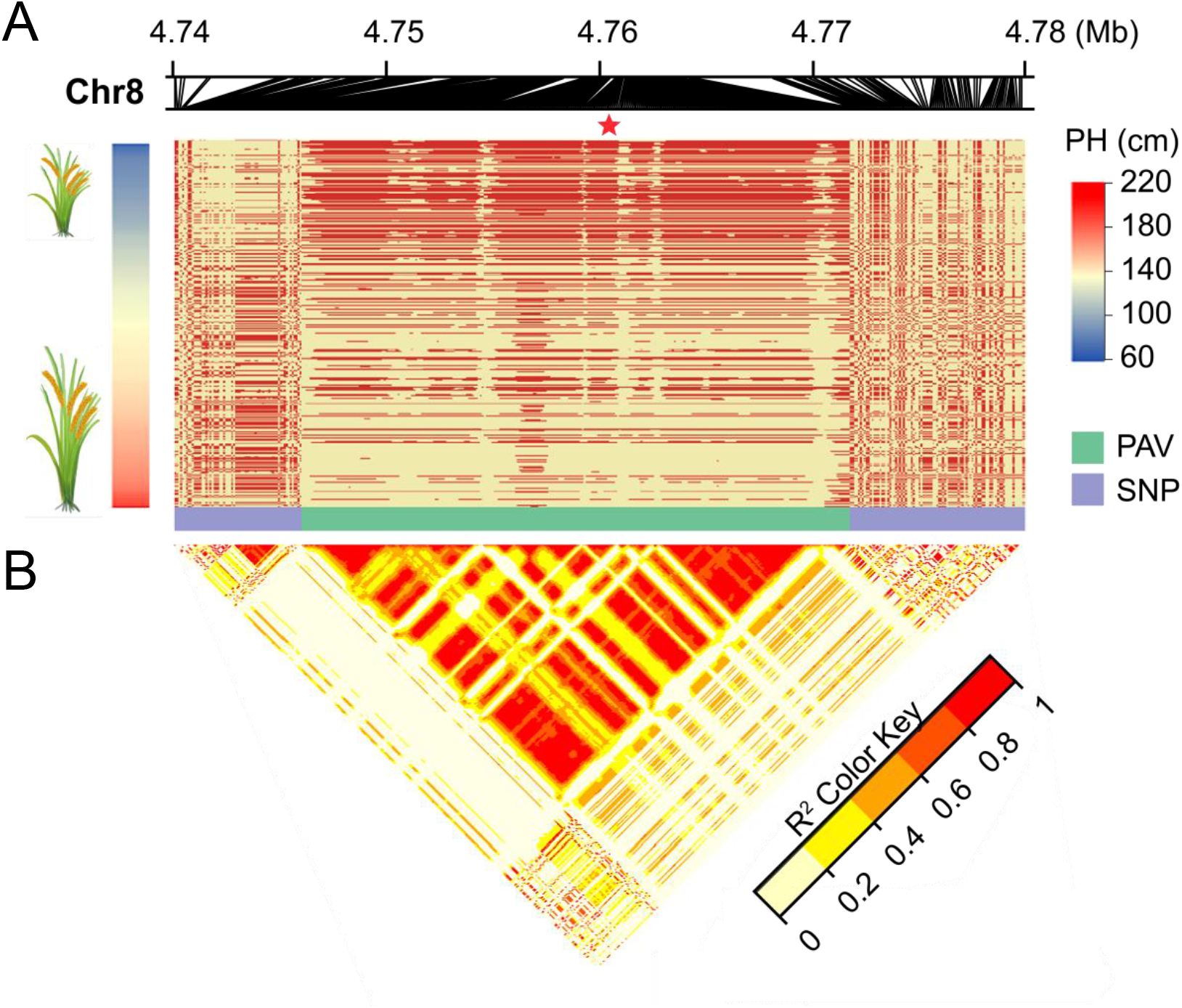
The relationship of PAVs and SNPs underlying plant height QTL in Chromosome 8. **A** Genotype of PAVs and SNPs display. In the PAV region, red bar means present of the PAV, yellow bar represents absence of the PAV. The five-pointed star indicates the position of the peak association PAV. **B** LD heatmap shows the regions surrounding the strong peaks of the PAV.

## Discussion

### PSVCP provides an accurate and robust tool for pangenome analysis

Many genomics studies include mapping sequencing data to reference genomes to identify genomic variation. However, these analyses suffer from bias due to the use of a single reference genome. Reference bias is especially problematic in the analysis of SVs, which is a major form of genomic variation in plants [29]. As an alternative, a pangenome can represent the genomic diversity of a species or population better than a single reference. Using a pangenome as a reference for mapping sequencing data supports accurate downstream analysis and avoids reference bias.

Currently, the most advanced method for pangenome construction and analysis is the graph-based strategy, which maintains the position of variable genetic information for each accession [14-16]. However, the graph-based pangenome approach also leads to challenges. This strategy is still in the early development stage, and plants lack a standard approach for graph-based pangenome construction and analysis. Furthermore, which are common in plants. Many pangenomic approaches stem from research on the human genome, which has much smaller genome variations between individuals than plant genomes. So graph-based pangenomes sometimes may not be able to fully represent large structural variations [30]. Furthermore, since plants contain complex repeat regions, they require significant computational resources for graph-based pangenome construction, especially for crops with large genome sizes. There are still insufficient tools available for the analysis of graph-based pangenomes. For example, while pangenome mapping algorithms have been developed for mapping reads to sequence graphs [31], none have challenged the dominance of linear genome-based mapping tools.

Because of the challenges in applying graph-based pangenomes, the linear pangenome is still useful for both functional genomic studies and breeding applications. In this study, we developed a new pipeline for constructing linear pangenomes (PSVCP) and aimed to overcome the bottleneck of other linear pangenome strategies. A major challenge for current linear pangenome construction strategies is the ability to accurately embed the newly identified PAV sequences into the linear reference. In several recent pangenome studies, including the 3,010 rice pangenome [12], the tomato pangenome [32] and *Brassica napus* pangenomes [4], novel sequences are placed as contigs that do not consider their genomic context. This limitation can limit further use of the pangenome in downstream gene mapping or functional validation of the candidate PAVs, since the nearby sequences may be important for the functional analysis of the PAVs. For example, a Pan-SV analysis in tomatoes revealed that the majority of gene-associated SVs are in cis-regulatory regions, and many are associated with subtle changes in expression [33]. To address this issue, PSVCP is designed to place novel sequences into the correct genome position, providing an accurate genetic map for functional genomic studies. The accuracy of the placement of the novel sequences by PSVCP was confirmed by successfully identifying the existence of the novel sequences and the sequence surrounding them by PCR amplification followed by sequencing. The advantage of our strategy was further demonstrated by GWAS analysis using PAV genotypes from our pangenome. Our PAV-GWAS successfully captured the casual structural variants of TGW and PH, while these variants are not available in the Nipponbare reference, or hard to characterize their biological meaning without the sequence information surrounding them. The pangenome constructed using PSVCP benefits from its linear format, which can directly integrate with currently available bioinformatics pipelines such as GATK [34] for genome variant discovery, and JBrowse [35] for genome visualization.

### PAVs provide insights into rice population structure

Most population structure studies are currently performed using SNPs [36], however, structural variants such as PAVs are increasingly used since they provide additional information about the population structure [4, 16, 32]. SV-based population structure studies are likely to become a tool for improving our understanding of the adaptation and evolution of species.

The rice pangenome constructed in this study contains novel genome sequences and annotated genes from comprehensive comparative genomic analysis. Our results indicate that compared to SNPs, PAVs provided further insights into rice evolution when used to identify genetic differentiation regions using Fst and phylogenetic inferences. In most cases, we found that SNP and PAV-based population structure analyses shared a similar Fst value change. However, in some genetic regions, PAV-based analysis has significant different Fst values than SNP-based results, providing higher resolution to differentiate the population structure. A 1.6 Mb interval in chromosome 8 displayed a much higher Fst value in PAV-based analysis than SNP-based analysis between *Japonica* and *Indica*. Higher frequencies of novel sequence insertions were discovered, which may be due to transposon movement in this region. More haplotype diversity was observed using PAVs than SNPs, suggesting that SNPs may underestimate genetic differentiation in some highly diverse genomic regions. These results demonstrate that PAV genotypes in our pangenome can provide additional power and information in analyzing genomic divergence and evolution.

Our results indicate that the majority of the newly inserted PAV sequences are transposable elements. Compared with SNP-based phylogeny, PAV-based phylogeny shows that some *Indica* accessions clustered with the *Aus* subpopulation, which is consistent with the TE-insertion phylogeny analysis using 3000 rice accessions [12]. This result also reflects the fact that *Aus* and *Indica* contain more common TE-insertions, since the divergence of the *Indica/Aus* lineages occurred more recently (∼540,000 years ago) than the divergence of *Japonica* (∼800,000 years ago). Additionally, introgression is potentially detected between *Indica* and *Aus* subpopulations based on the PAV data, consistent with previous studies showing that *Indica* accessions contain *Aus* introgressions [37] and *Indica* and *Aus* show closer genetic affinity [38]. The phylogeny variations between SNP and PAV analysis are consistent with observations in other plants such as *Arabidopsis* [39], *Amborella trichopoda* [25], green millet *Setaria viridis* [40] and *Brassica oleracea* [4], showing that PAV or SV can provide additional information to characterize population structure that might associate with transposon movement during genome evolution, highlighting the value of using PAVs or SVs in addition to SNPs in assessing species evolution.

### PAV-based GWAS provides additional power to identify causal variants

Most GWAS analysis uses SNPs identified from a single reference genome as markers to detect marker-trait associations. However, recent studies suggest that SVs, including PAVs, contribute to and explain more variation than SNPs for many traits [41]. Phenotypes associated with regions that are absent in the reference genome can only be mapped to a region in the LD block linked with the PAV. However, this association cannot be identified if the PAV haplotypes are not in LD with the SNPs surrounding them, which we observed in our results (Fig. 7A). Furthermore, using variation identified from a single reference in GWAS may cause bias, which weakens the ability of GWAS to identify associations. For example, a maize gene conferring resistance to sugarcane mosaic virus is present in the B73 reference but not in the PH207 reference. Conducting GWAS using SNPs genotyped using the B73 reference can identify the gene, while the PH207 cannot [42]. Using PAVs identified from a pangenome can help resolve the above problems, and PAVs can complement SNP-based GWAS. For example, a recent study in *Brassica napus* shows that a PAV-based pangenome-wide association study can directly pinpoint the causal SVs for silique length, seed weight and flowering time [43].

In this study, PAVs are genotyped from the pangenome constructed by the PSVCP pipeline, and used for GWAS analysis of TGW and PH in an international rice panel. Both PAV-GWAS and SNP-GWAS methods can identify previous characterized QTLs, such as *GW5* for TGW and *sd1* for PH. Surprisingly, the peak PAV-GWAS signals are directly and accurately located in the functional PAVs, causing the phenotypic variations, while the most significant signal for SNP-GWAS can only identify the approximate location of the causal variants.

Importantly, PAV-GWAS can identify new candidate causal variations that SNP-GWAS cannot discover. In our study, a 13 kb PAV containing two retrotransposons was found to be strongly associated with plant height using PAV-GWAS, and this was not identified using SNP-GWAS. Transposon movements are important sources of phenotypic variants. A GWAS study in tomatoes based on TE insertion polymorphisms revealed that transposon movement was associated with leaf morphology and fruit colour [44]. Further investigation of the 13-kb rice PAV showed that it was 2 kb upstream from *LOC_Os08g07400*, whose expression was associated with the present and absence of the 13-kb sequence. These results suggest that retrotransposon movement in this locus may lead to phenotypic variation by affecting the promoter region of LOC_Os08g07400.

To unravel why SNP-GWAS cannot identify this locus, we investigated the candidate variant region at a population level. Our results show that no SNPs were found in the 13 kb PAV sequence, and SNPs located near the 13 kb PAV sequence show a poor correlation with the PAVs, with no association between SNPs and the plant height phenotype. TEs having a low LD with nearby SNPs were observed in other genomic studies in rice and tomato [45]. Akakpo et al. (2020) reported that TE-GWAS could identify a signal associated with rice grain width on chromosome 4 that was missing in SNP-GWAS [46]. Recent retrotransposon insertion may cause the low LD of SNPs by breaking previous linkage disequilibrium. However, further investigation is required to understand how they affect functional gene expression and phenotype variation. Our study demonstrates that a PAV-based pangenome-wide association analysis is a powerful approach to detect and dissect the genetic variants causing phenotypic variation of agronomical traits.

### Conclusions

A new strategy and pipeline to construct a linear pangenome by whole genome comparison were developed in the present study. This strategy supported the construction of a linear pangenome that can solve the problems of preserving the location information of SVs and facilitates downstream pangenomic analysis. A rice pangenome was constructed using 12 complete genomes spanning all rice subpopulations. Downstream population analysis demonstrated that using the pangenome provided insights into the rice population structure and evolution, which are not available by analysis using SNPs from a single reference. GWAS analysis using the pangenome reference revealed a significant improvement in power, especially in characterizing causal PAVs. The new pangenome construction pipeline and the rice pangenome provide a novel framework for future pangenomic studies in rice and other plants.

## Methods

### Plant materials

Seed for 413 accessions was sown on July 28th, 2020, at Guangzhou, Guangdong, China. High-molecular-weight genomic DNA was extracted from 30-day-old leaves following a standard CTAB (hexadecyltrimethylammonium bromide) protocol. Sequencing was performed on the Illumina NovaSeq6000 platform (BerryGenomics, China). A fastx_toolkit (http://hannonlab.cshl.edu/fastx_toolkit) was used to remove adaptor and low-quality reads. All reads have been deposited in the NCBI sequence read archive (BioProject accession PRJNA820969). Plant height and thousand grain weight were assessed at the mature growth stage with three biological replicates.

### Construction of the pangenome

Data for twelve assembled genomes were downloaded from the Rice Resource Center (https://ricerc.sicau.edu.cn/) [10], representing MSU, Lemont, NamRoo, LJ, CN1, R498, TM, Tumba, FH838, N22, Basmati1 and CG14. We employed an iterative strategy to construct the pangenome. First, we carried out pairwise collinearity comparison between NIP and Lemont using MUMmer 4.0.0 [47], with parameters: ‘‘--maxgap 500 --mincluster 1000 --diagdiff 20”. NIP was named as ref0. We used Assemblytics to detect and analyze variants from MUMmer. SVs were identified by comparison of the first genome (Lemont) with the Nipponbare reference genome assembly (ref0). The insertions larger than 50 bp were identified and incorporated to generate the new reference genome (ref1). The ref1 genome was then further compared with each genome iteratively until all genomes were incorporated into the pangenome (Additional file 2: Table S1).

### Short read data processing for PAV-GWAS

Paired-end short-read sequencing data for each accession was aligned to the pangenome using BWA-MEM [48]. Mapping results were sorted using Picard and filtered using SAMtools [49], retaining reads with a mapping quality over 20. We used the SAMtools with the parameters: ‘‘-F 4 -F 256” to remove reads that did not map to the pangenome or mapped to the pangenome repeatedly. Using the pangenome as the reference genome, the coverage of each accession was detected in every 20 bp region by Mosdepth [50] with the parameters: “-b 20”. Two adjacent 20 bp regions were merged if adjacent sequences had coverage of >5 reads.

### PAV identification

PAVs were called based on the coverage for each accession. We combined all PAV information by row into a map, displayed as a matrix (Fig 1C) with accession names as rows. Segments were defined as PAV regions, named by the adjacent left breakpoint position, and the population PAV genotype matrix was filtered by minor allele frequency (MAF) >0.05.

### Gene PAV detection

A gene was considered missing when the horizontal coverage across the CDS is less than 95% and the vertical coverage less than two, as used in the 3K-RG study [12] using Mosdepth v0.2.6 [50]. A PAV matrix was generated showing the presence or absence of each gene for each accession. The statistical significance of gene frequency changes was calculated using Fisher’s exact test. P-values were adjusted for multiple comparisons using the Bonferroni method as implemented in p.adjust from R v3.5.0. Genes with an adjusted p-value<0.001 and difference frequency between groups ≥10%[32] were defined as significant.

### Short read data processing for SNP-GWAS

Short read resequencing data were aligned to the NIP reference genome using BWA-MEM. The results were sorted using Picard and filtered using SAMtools, retaining reads with a mapping quality over 20. Nucleotide variants for each accession were detected using HaplotypeCaller in GATK (v3.8-1-0) [34] with the default parameters. Population nucleotide variants were called using CombineGVCFs and GenotypeGVCFs tool in GATK. Finally, we used the SelectVariants and VariantFiltration tool in GATK to filter the genotype of the population.

### GWAS analysis

To construct the PAV genotype map for GWAS, we used “A” representing “Absent” and “C” to represent “Present” in the HapMap genotype file. PAVs and SNPs were selected for GWAS analysis based on the criteria of missing data <15% and minor allele frequency of >0.05. GWAS was performed using a mixed linear model (MLM) with kinship matrix and principal component analysis in GAPIT version 2 [51]. The significance cutoff was defined as the threshold of –log_10_(*p*) <5. Manhattan plots were produced using CMplot package (https://github.com/YinLiLin/R-CMplot) in R v3.5.0.

### GO analysis

Functional annotation was performed using Blast2GO v2.5 [52]. Genes were aligned to the proteins in the Viridiplantae database using BLASTP [53] (E-values <1 × 10^−5^). Gene ontology (GO) analysis was conducted using topGO [54] and Fisher’s exact test with ‘elim’ was used to correct for multiple comparisons.

### Population structure and genotype analysis

Filtered PAV and SNP data were used for the population structure study. SNP-based and PAV-based phylogenetic trees of 413 rice accessions were constructed using IQ-tree using a maximum likelihood method (with the alrt 1000 -bb 1000), respectively. SNP-based and PAV-based principal component analyses were performed with GCTA (Genome-wide Complex Trait Analysis) v1.93.2 [55]. SNP-based and PAV-based Fixation index (Fst) values were calculated using a 100 kb sliding window (with a 10 kb step for FST values calculation) using VCFtools [56].

### TE analysis

A *de novo* transposable element (TE) library was generated for the rice pangenome using EDTA v1. Using BLAST+ v 2.2.3 [53], the representative retrotransposon TE families in Carpentier et al. (2019) [22] were used to search the rice pangenome library to identify the whole genome-wide TEs (with >85% sequence identity and e-value < 10^−5^).

## Supplementary Information

Additional file 1. Supplemental figures 1-9

Additional file 2. Supplemental tables 1-11

## Authors’ contributions

JW, WY, HH, CL, DE and JZ designed the research. JW, WY, HH and JZ conducted the experiments and analyzed the data. JW, WY, YM, HH, CL, DE and JZ wrote the original draft and edited the manuscript. Other authors assisted in experiments and discussed the results. All authors read and approved the final manuscript.

## Acknowledgement

The sutdy was supported by Guangdong Provincial International Cooperation Project of Science & Technology (2021A0505030048), the Innovation Team Project of Guangdong Modern Agricultural Industrial System (2022KJ106), the “YouGu” Plan of Rice Research Institute of Guangdong Academy of Agricultural Sciences (2021YG001), the Evaluation and Operation Funds of Guangdong Key Laboratories (2020B1212060047), Special Fund for Scientific Innovation Strategy-Construction of High Level Academy of Agriculture Science (R2019PY-JX001, R2019YJ-YB3006). the GuangDong Basic and Applied Basic Research Foundation (2020A1515111025). The authors would also like to thank the Pawsey Supercomputing Centre for the use of their computing resources.

## Availability of data and materials

The raw read data (FASTQ files) of 413 accessions were uploaded to NCBI’s sequence read archive (BioProject accession PRJNA820969). PSVCP is freely available at (https://github.com/wjian8/psvcp_v1.01).

