## Supplementary Figures for "A pangenome analysis pipeline (PSVCP) provides insights into rice functional gene identification"

### Slide 1
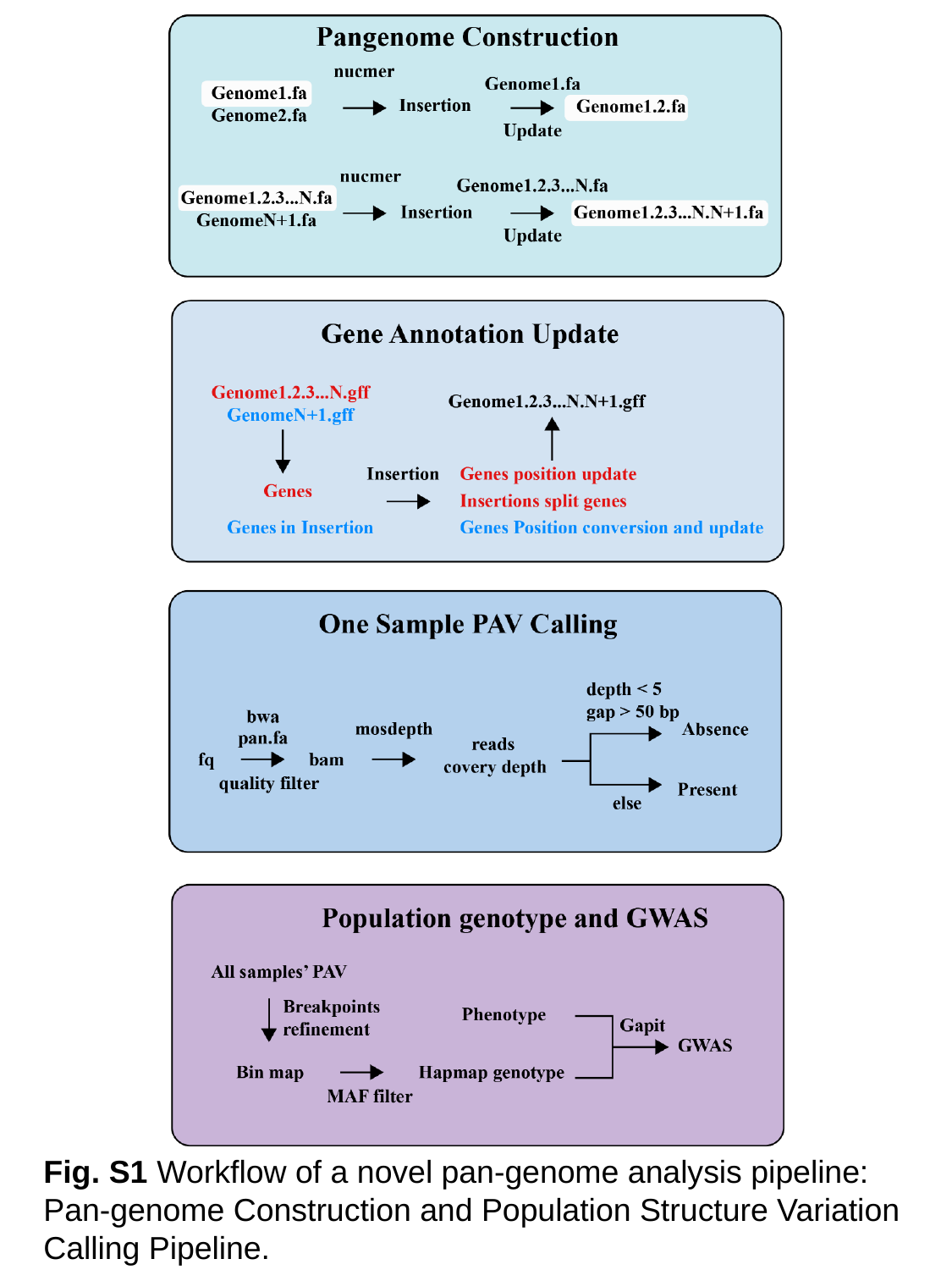

Fig. S1 Workflow of a novel pan-genome analysis pipeline: Pan-genome Construction and Population Structure Variation Calling Pipeline.

### Slide 2
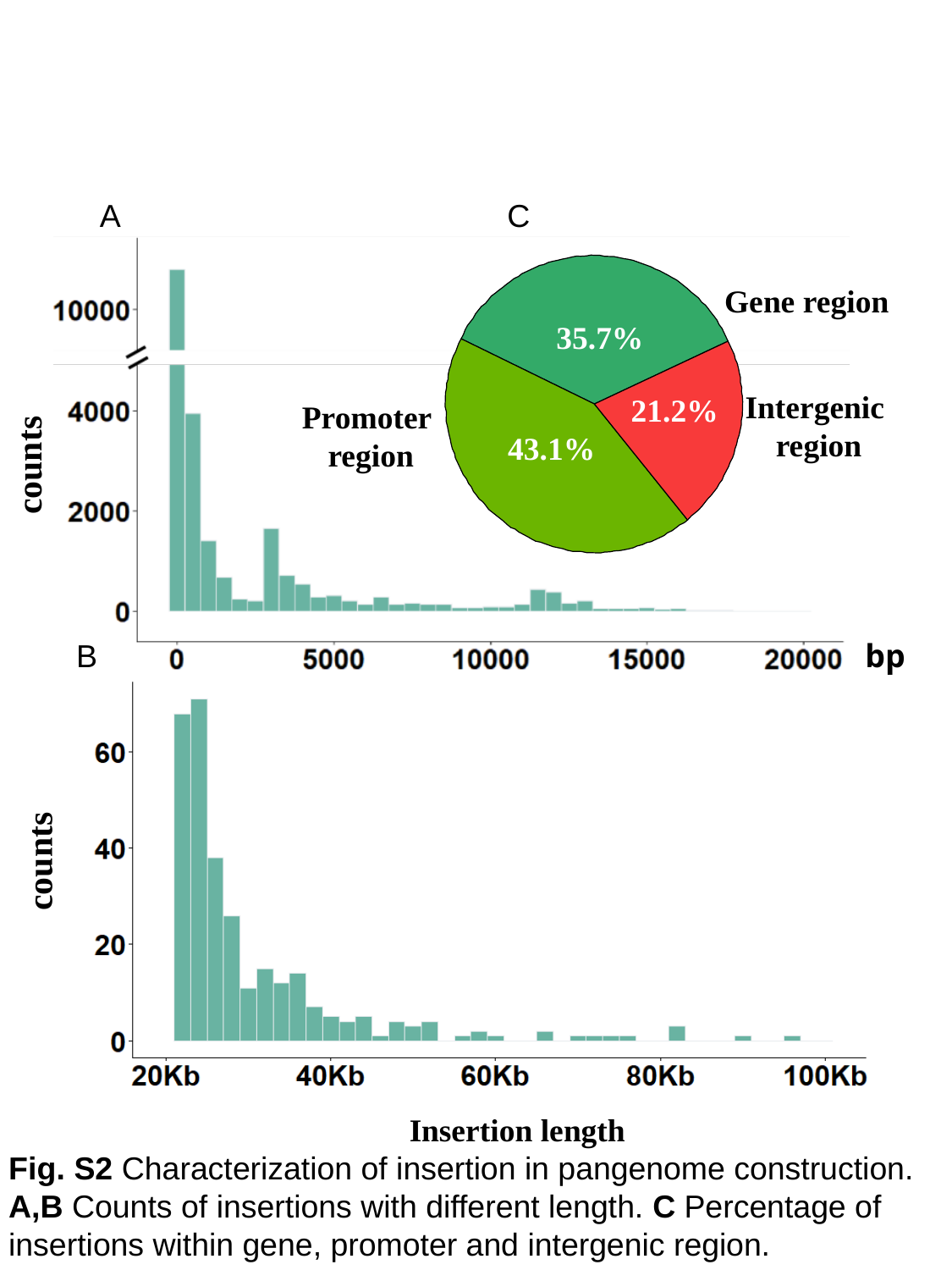

A
C
counts
bp
counts
Insertion length
35.7%
21.2%
43.1%
Gene region
Intergenic
region
Promoter
region
B
Fig. S2 Characterization of insertion in pangenome construction. A,B Counts of insertions with different length. C Percentage of insertions within gene, promoter and intergenic region.

### Slide 3
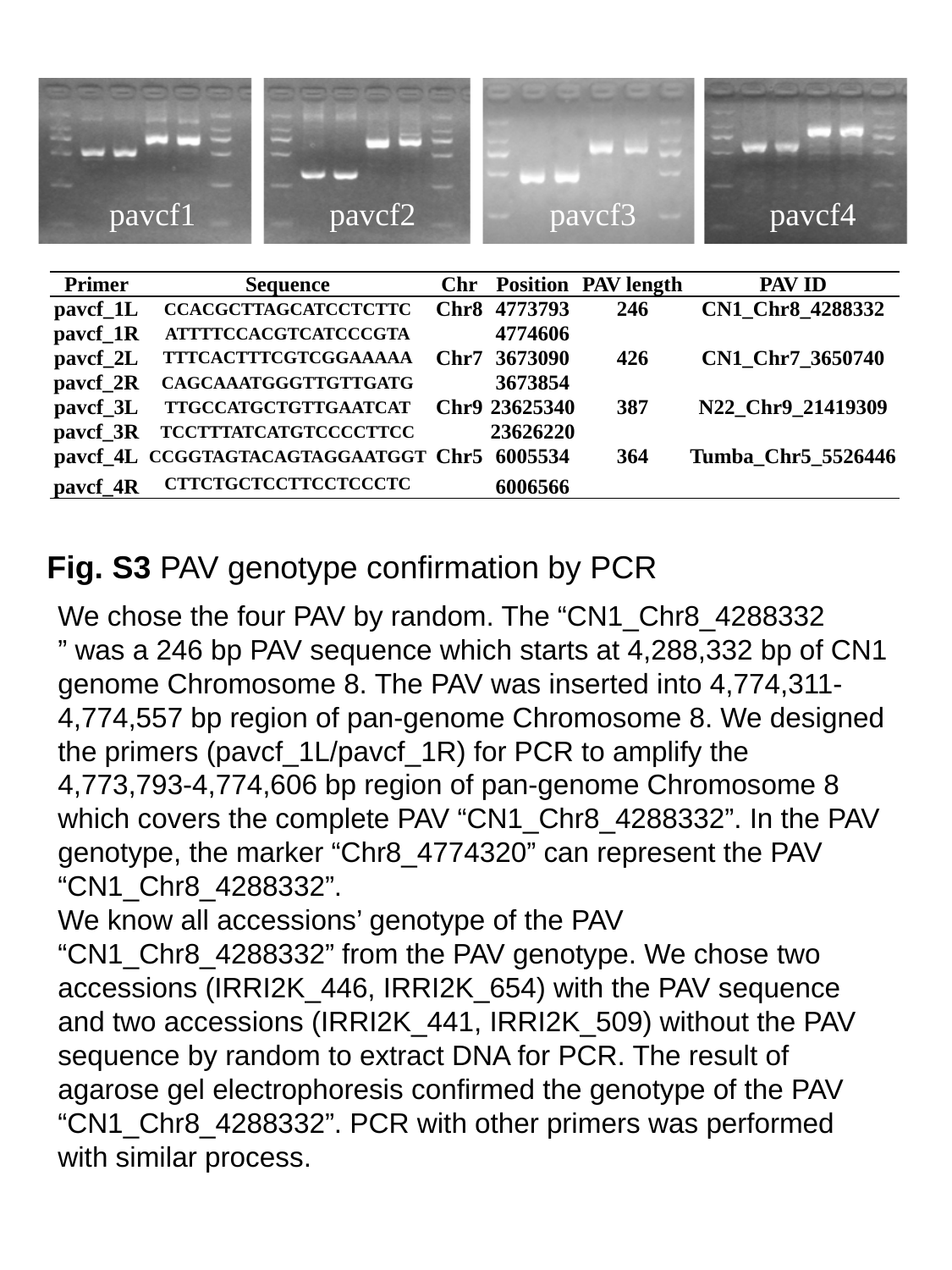

pavcf1
pavcf2
pavcf3
pavcf4
| Primer | Sequence | Chr | Position | PAV length | PAV ID |
| --- | --- | --- | --- | --- | --- |
| pavcf\_1L | CCACGCTTAGCATCCTCTTC | Chr8 | 4773793 | 246 | CN1\_Chr8\_4288332 |
| pavcf\_1R | ATTTTCCACGTCATCCCGTA | | 4774606 | | |
| pavcf\_2L | TTTCACTTTCGTCGGAAAAA | Chr7 | 3673090 | 426 | CN1\_Chr7\_3650740 |
| pavcf\_2R | CAGCAAATGGGTTGTTGATG | | 3673854 | | |
| pavcf\_3L | TTGCCATGCTGTTGAATCAT | Chr9 | 23625340 | 387 | N22\_Chr9\_21419309 |
| pavcf\_3R | TCCTTTATCATGTCCCCTTCC | | 23626220 | | |
| pavcf\_4L | CCGGTAGTACAGTAGGAATGGT | Chr5 | 6005534 | 364 | Tumba\_Chr5\_5526446 |
| pavcf\_4R | CTTCTGCTCCTTCCTCCCTC | | 6006566 | | |
Fig. S3 PAV genotype confirmation by PCR
We chose the four PAV by random. The “CN1_Chr8_4288332
” was a 246 bp PAV sequence which starts at 4,288,332 bp of CN1 genome Chromosome 8. The PAV was inserted into 4,774,311-4,774,557 bp region of pan-genome Chromosome 8. We designed the primers (pavcf_1L/pavcf_1R) for PCR to amplify the 4,773,793-4,774,606 bp region of pan-genome Chromosome 8 which covers the complete PAV “CN1_Chr8_4288332”. In the PAV genotype, the marker “Chr8_4774320” can represent the PAV “CN1_Chr8_4288332”.
We know all accessions’ genotype of the PAV “CN1_Chr8_4288332” from the PAV genotype. We chose two accessions (IRRI2K_446, IRRI2K_654) with the PAV sequence and two accessions (IRRI2K_441, IRRI2K_509) without the PAV sequence by random to extract DNA for PCR. The result of agarose gel electrophoresis confirmed the genotype of the PAV “CN1_Chr8_4288332”. PCR with other primers was performed with similar process.

### Slide 4
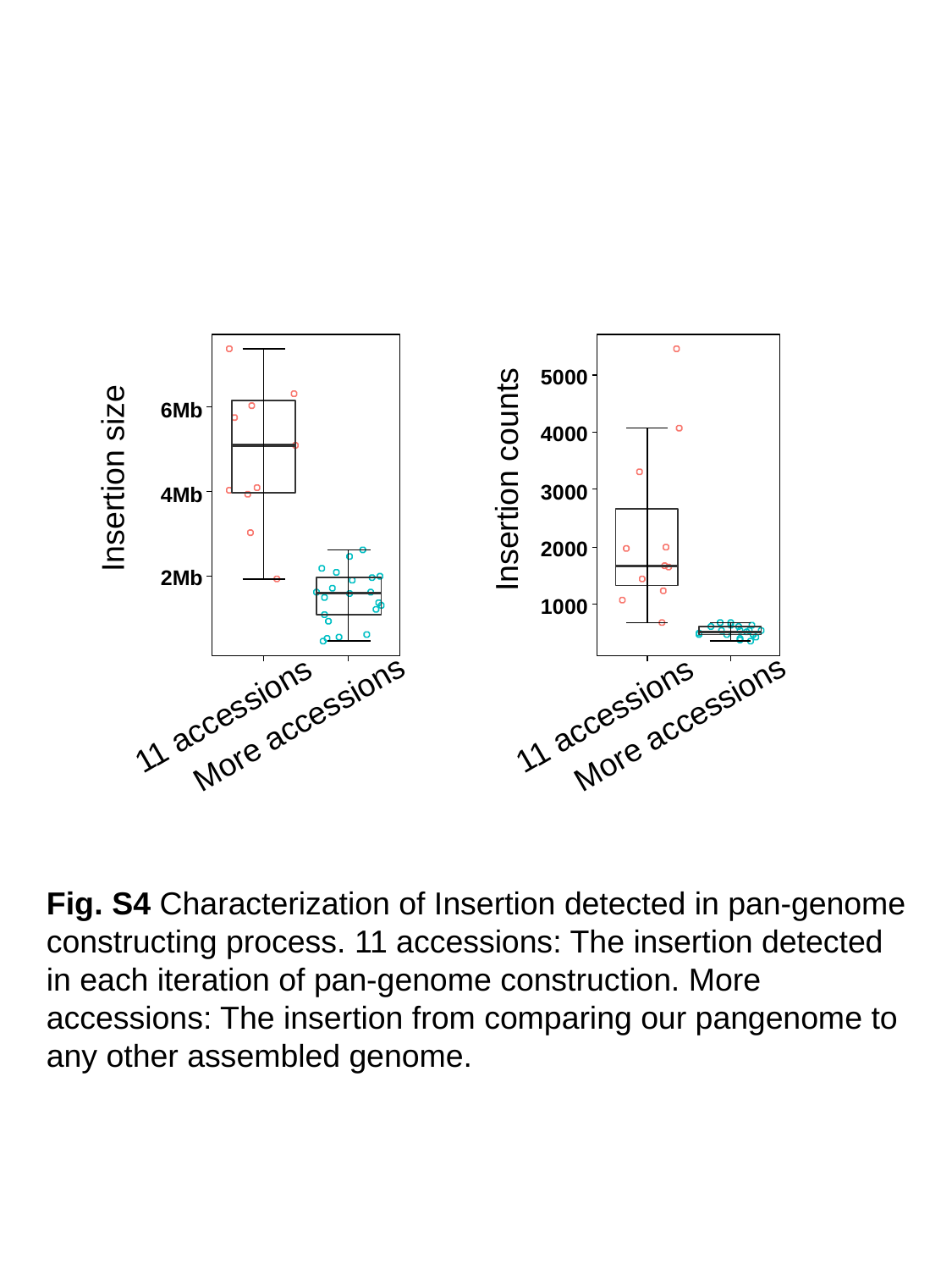

Insertion size
Insertion counts
11 accessions
11 accessions
More accessions
More accessions
Fig. S4 Characterization of Insertion detected in pan-genome constructing process. 11 accessions: The insertion detected in each iteration of pan-genome construction. More accessions: The insertion from comparing our pangenome to any other assembled genome.

### Slide 5
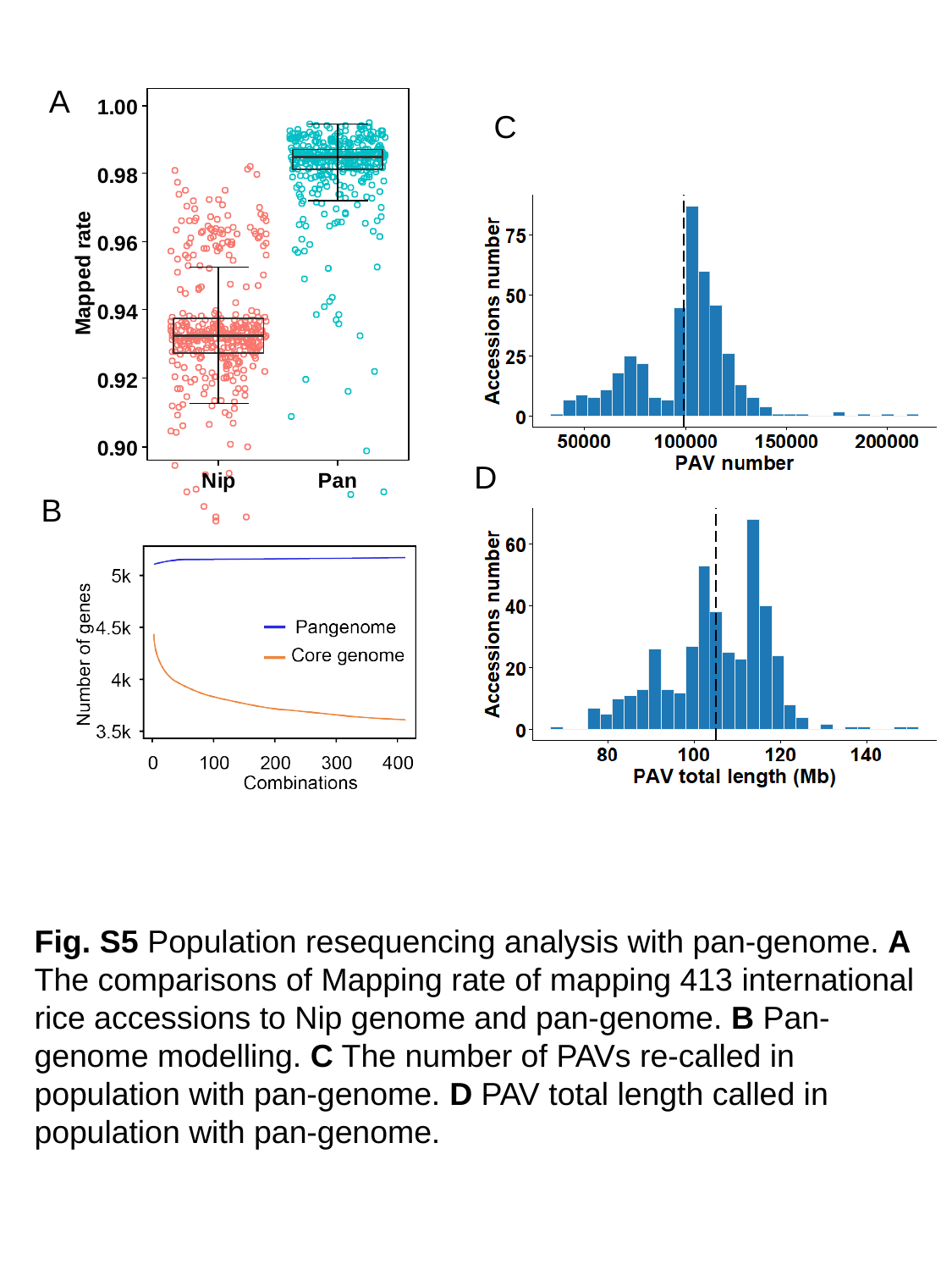

A
C
D
B
Fig. S5 Population resequencing analysis with pan-genome. A
The comparisons of Mapping rate of mapping 413 international rice accessions to Nip genome and pan-genome. B Pan-genome modelling. C The number of PAVs re-called in population with pan-genome. D PAV total length called in population with pan-genome.

### Slide 6
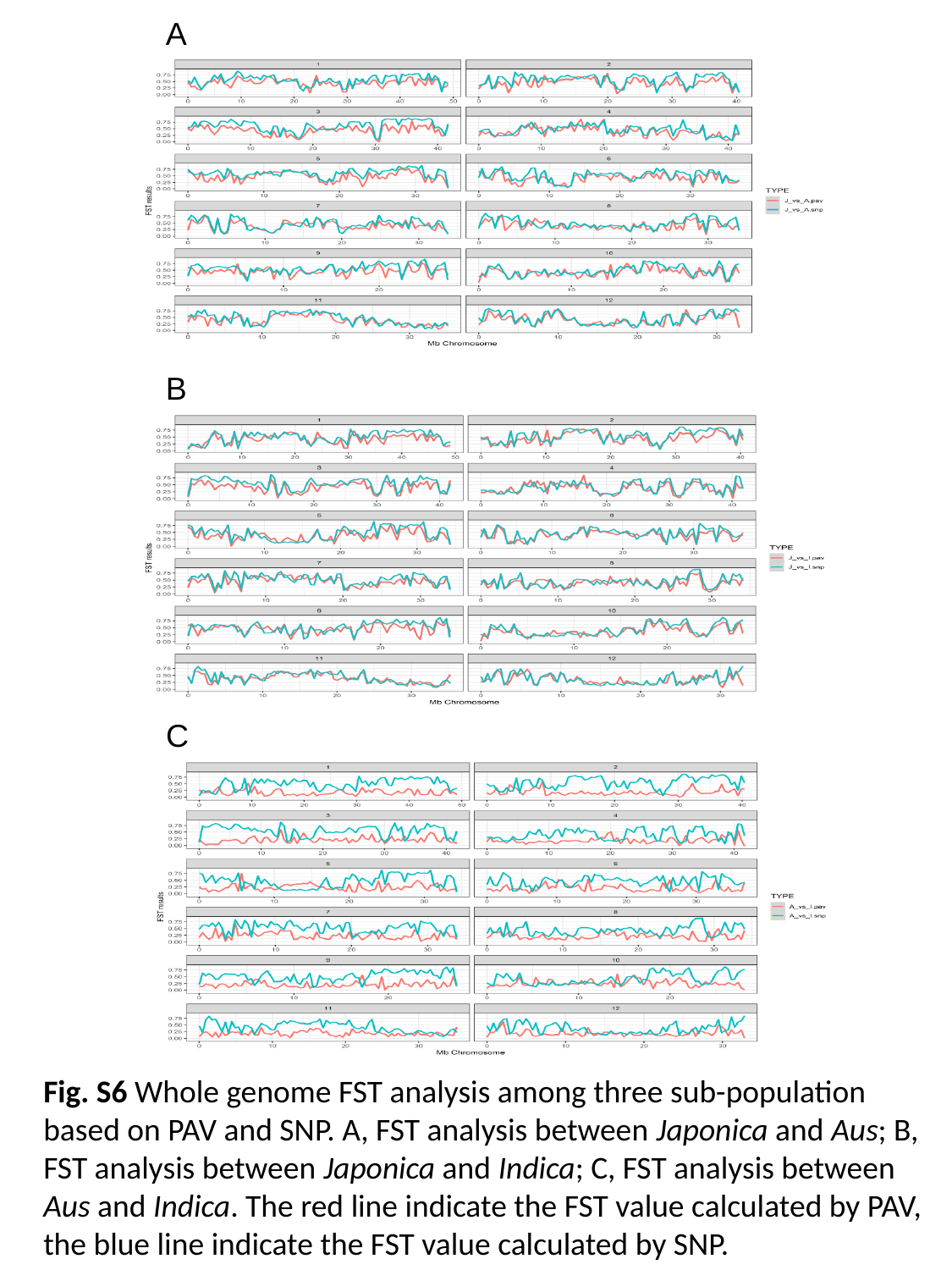

A
B
C
Fig. S6 Whole genome FST analysis among three sub-population based on PAV and SNP. A, FST analysis between Japonica and Aus; B, FST analysis between Japonica and Indica; C, FST analysis between Aus and Indica. The red line indicate the FST value calculated by PAV, the blue line indicate the FST value calculated by SNP.

### Slide 7
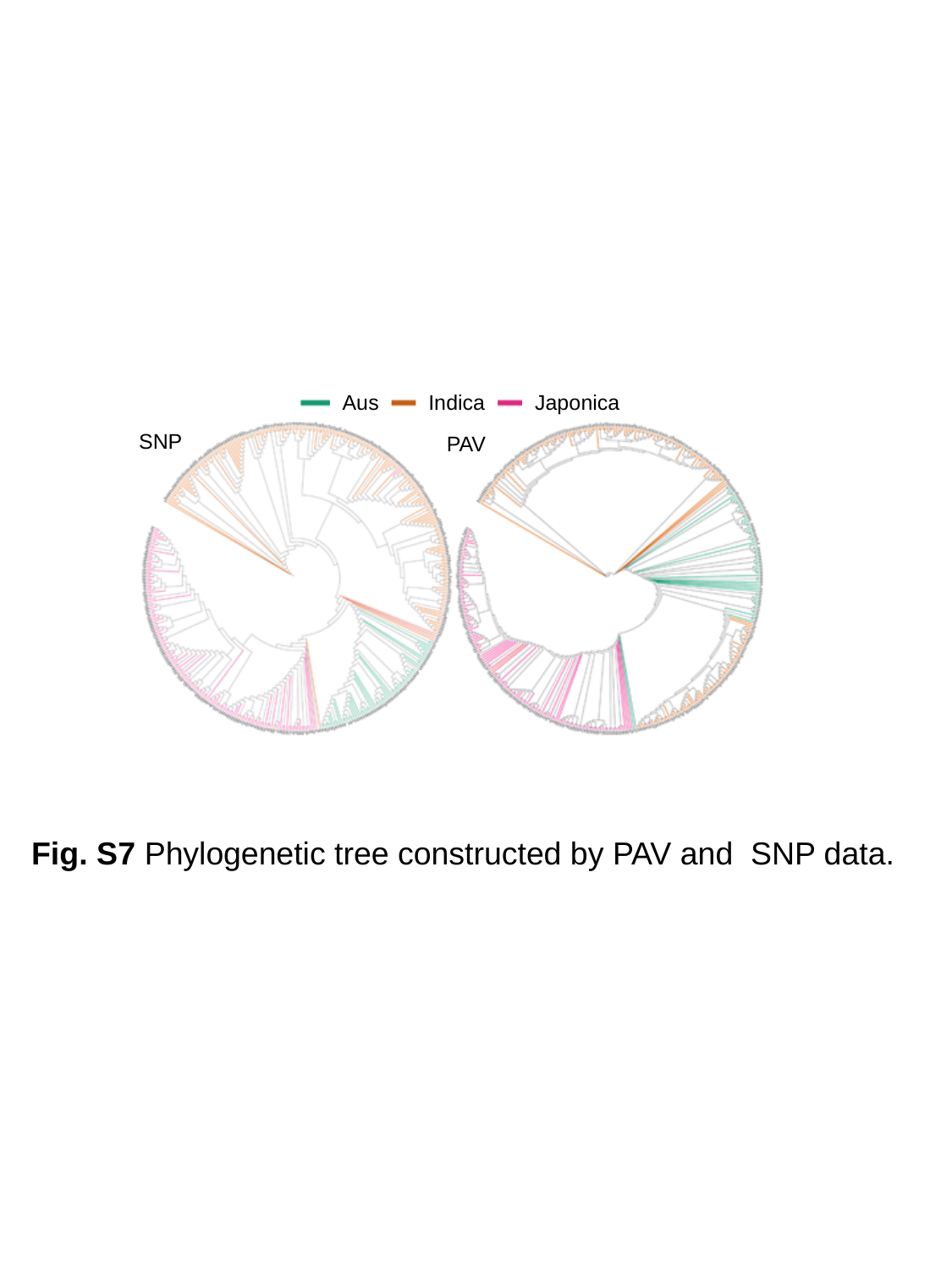

Japonica
Aus
Indica
SNP
PAV
Fig. S7 Phylogenetic tree constructed by PAV and SNP data.

### Slide 8
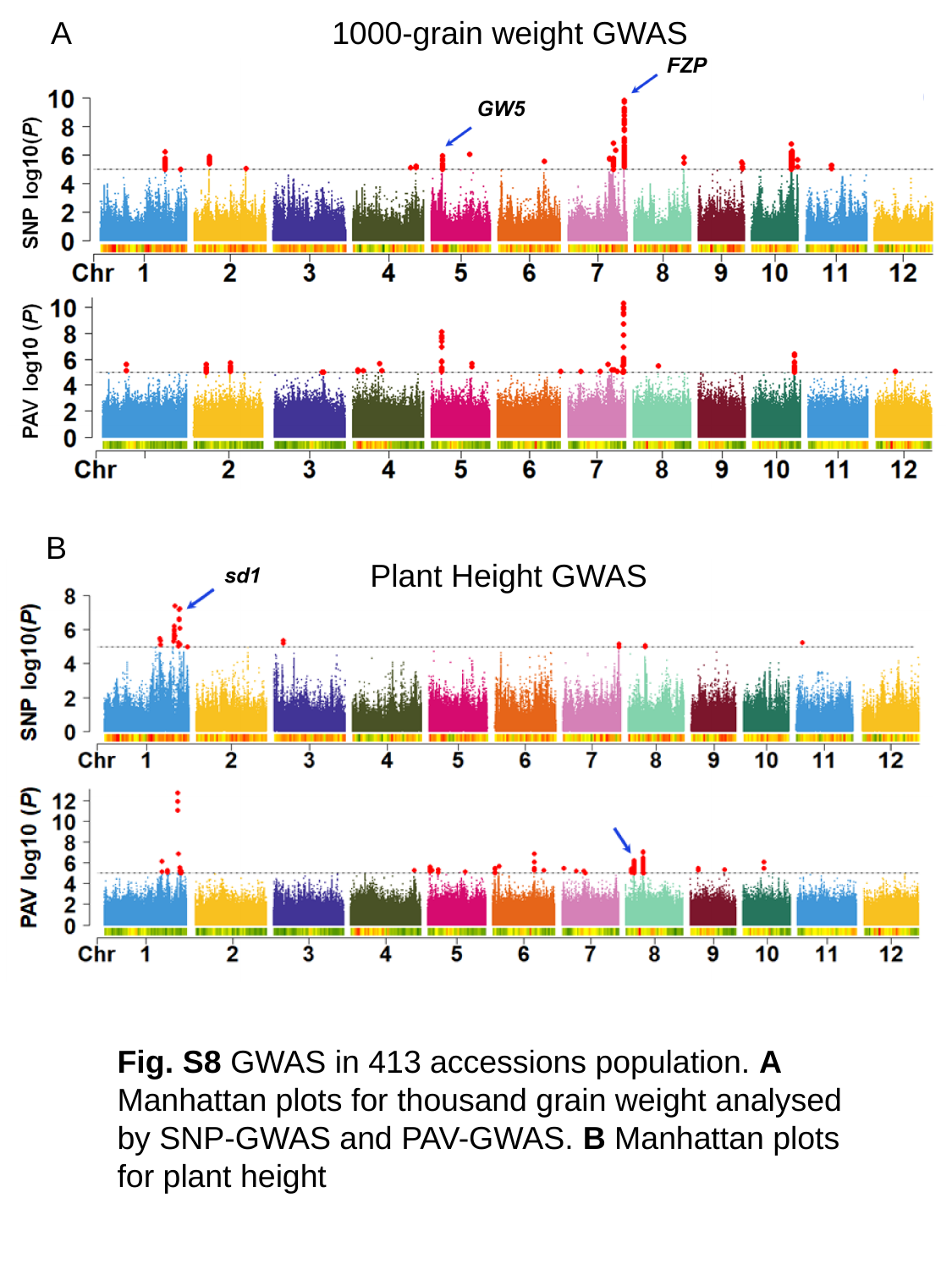

A
1000-grain weight GWAS
B
Plant Height GWAS
Fig. S8 GWAS in 413 accessions population. A Manhattan plots for thousand grain weight analysed by SNP-GWAS and PAV-GWAS. B Manhattan plots for plant height

### Slide 9
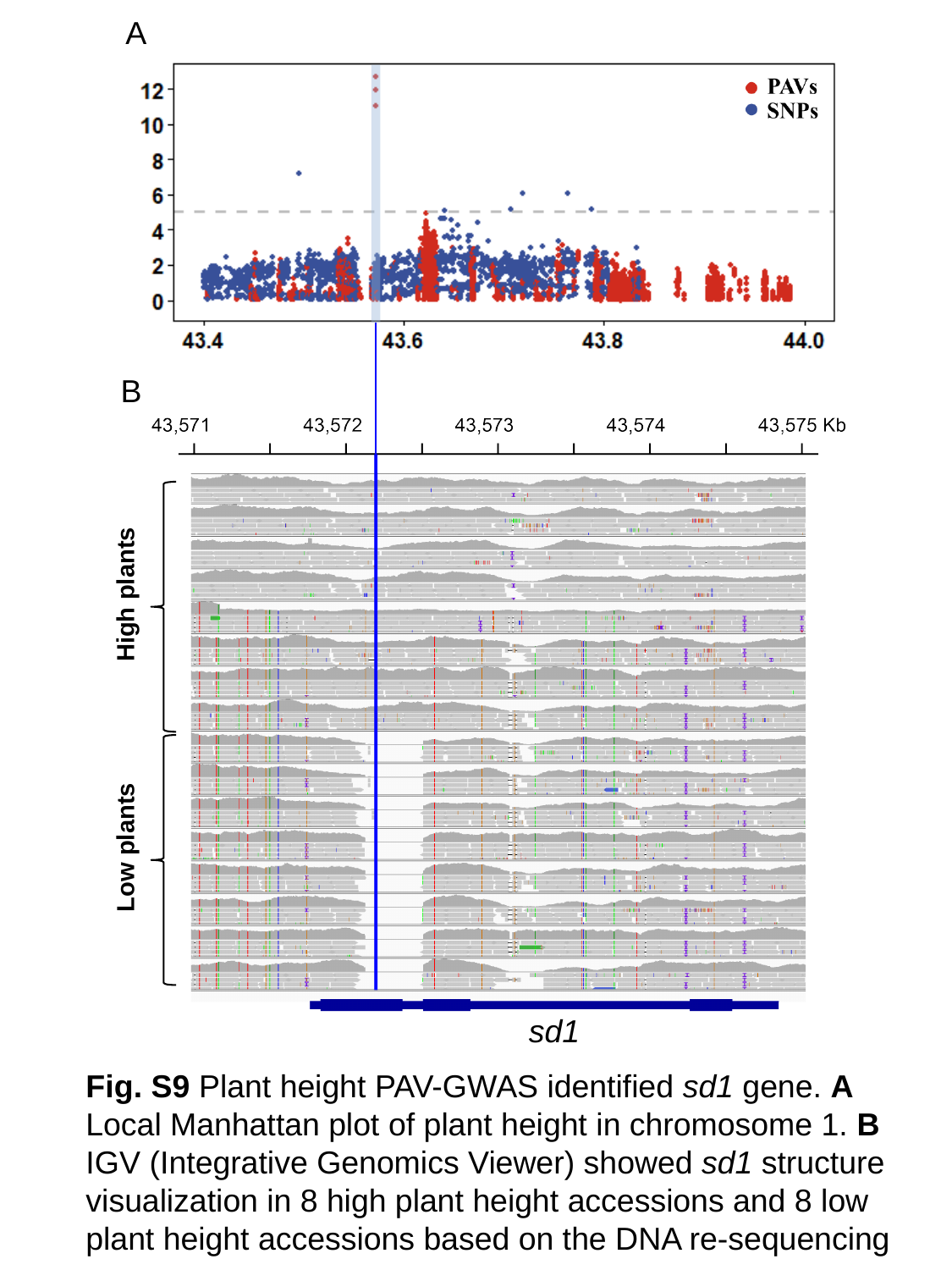

A
B
High plants
Low plants
sd1
Fig. S9 Plant height PAV-GWAS identified sd1 gene. A Local Manhattan plot of plant height in chromosome 1. B IGV (Integrative Genomics Viewer) showed sd1 structure visualization in 8 high plant height accessions and 8 low plant height accessions based on the DNA re-sequencing
